# MALDI-ToF detection of *Leishmania infantum* infection in *Lutzomyia longipalpis* and *Nyssomyia neivai*

**DOI:** 10.64898/2026.06.18.733076

**Authors:** Lucas Alexandre Farias de Souza, Emilie Kariya, Jorian Prudhomme, Jérôme Depaquit, Magda Clara Vieira da Costa-Ribeiro, Antoine Huguenin

**Author notes:** **Corresponding author**: Laboratoire de Parasitologie-Mycologie, pôle de Biopathologie, CHU de Reims, 51097 Reims, France. mail.

## Abstract

**Background:** Matrix-assisted laser desorption/ionization time-of-flight mass spectrometry (MALDI-ToF MS) is widely used for sand fly identification, but its potential to detect *Leishmania* infections in vectors remain underexplored. This pilot study evaluated whether MALDI-ToF MS protein profiles of lab-reared *Lutzomyia longipalpis* and *Nyssomyia neivai* can discriminate *Leishmania infantum*–infected from uninfected females.

**Methodology:** Colonies were experimentally infected with *L. infantum* using membrane feeding, and females were collected at different days post-blood meal. Thoraces and legs were processed individually for MALDI-ToF MS, and spectra were analysed using both Bruker software and custom R pipelines.

**Principal findings:** Unsupervised approaches (MSP dendrograms, PCA) showed limited or inconsistent separation of infection status for *Lu. longipalpis*. In contrast, supervised machine-learning models built on peak-intensity matrices achieved excellent discrimination between infected and uninfected specimens for both species, with several algorithms reaching near-perfect performance on an external test set not used for training. Variable-importance analysis highlighted sets of m/z peaks, mainly showing decreased intensity in infected sand flies, as putative infection biomarkers.

**Conclusion:** This proof-of-concept study highlights that *L. infantum* infection induces reproducible, species-specific alterations in sand-fly MALDI-TOF profiles, supporting further development of high-throughput, MS-based screening of infected vectors.

**Author summary:** *Leishmania infantum* is a parasite responsible for visceral leishmaniasis, a severe neglected tropical disease. It is transmitted to humans by sandfly vectors. This study explored whether the MALDI-ToF mass spectrometry technique can detect infection by the *L. infantum* parasite in the two main sandfly vectors in Brazil: *Lutzomyia longipalpis* and *Nyssomyia neivai*. The method has already been tested to identify sandfly species, but its ability to detect infected insects had not been well studied. We infected laboratory-reared sandflies and analyzed their protein profiles to see whether infected and uninfected individuals could be distinguished.

We found that infection changes the molecular fingerprints of both sandfly species. Machine-learning models were able to distinguish infected from uninfected specimens with very high accuracy. A small part of the most informative signal was shared between both species, while most of the peaks were species-specific, suggesting that infection affects each vector in a slightly different way.

These results show that MALDI-ToF has promise as a rapid, low-cost tool for screening sandflies for *Leishmania* infection. With further validation, this approach could complement existing surveillance methods and help monitor disease transmission in endemic areas.

## Introduction

Leishmaniasis comprises a group of parasitic diseases caused by protozoa of the genus *Leishmania* and transmitted by phlebotomine sand flies. It mainly occurs in two clinical forms: Cutaneous Leishmaniasis (CL) and Visceral Leishmaniasis (VL) [1]. These forms differ in clinical presentation, mortality, and socioeconomic impact, with VL being potentially fatal if untreated [2]. The disease is endemic in nearly 99 countries, disproportionately affecting populations in East Africa, South Asia, the Mediterranean basin, and Latin America [3,4]. The distribution of cases is highly heterogeneous, with a limited number of countries accounting for most cases. In Latin America, Brazil stands out as one of the countries with the highest number of CL cases and contributes substantially to VL cases across the region [5].

Among the more than 500 phlebotomine sand fly species described in the Americas [6], only a few are proven vectors of *Leishmania* . In Brazil, *Lutzomyia longipalpis* (Lutz & Neiva, 1912) is recognized as the principal vector of *Leishmania infantum*, the etiological agent of VL. This species exhibits high adaptability to urban environments and anthropogenic habitats, facilitating the maintenance of transmission cycles close to human dwellings [7].

On the other hand, *Nyssomyia neivai* (Pinto, 1926), traditionally associated with *L. braziliensis* transmission and CL, has been increasingly investigated for its potential role in *L. infantum* transmission. Although not yet recognized as a proven vector, recent molecular studies have detected *L. infantum* DNA in field-captured specimens of *Ny. neivai* in Southern Brazil [8,9].

These findings provide molecular evidence of *Ny. neivai*’s permissive susceptibility to *L. infantum*, especially in areas of absence of *Lu. longipalpis*, where *Ny. neivai* may act as a vector for the parasite. Such evidence highlights the need for integrated entomological surveillance and the use of advanced identification techniques, including Matrix Associated Laser Desorption-Ionization – Time Of Flight mass spectrometry (MALDI-ToF), to further clarify the epidemiological roles of both species in visceral leishmaniasis transmission cycles.

Vector surveillance is crucial, highlighting the need for a rapid and cost-effective tool to identify leishmaniasis vectors.

The MALDI-ToF MS has emerged as a reliable method for identifying sandflies, even in regions with high species diversity. Compared to traditional identification based on morphological criteria or molecular methods, this technique offers rapid, cost-effective species differentiation, which is crucial for the epidemiological surveillance of leishmaniasis vectors. This technique also makes the identification of sandflies accessible to non-specialists, compensating for part of the loss of entomological expertise.

A study conducted in French Guiana [10] demonstrated high accuracy using a custom reference database comprising 29 sand fly species. The database covered ecologically diverse Northern Amazonian species, including known *Leishmania* vectors. However, it did not include *Lu. longipalpis* species complex, key vectors of *L. infantum* the agent of visceral leishmaniasis in the New World. Moreover, *Nyssomyia* spp are only identified at the genus level.

MALDI-ToF has been used to discriminate between infected and non-infected vectors. It has been first used for detecting *Borrelia crocidurae* (a bacteria responsible for relapsing fever) in *Ornithodoros sonrai* soft ticks with 88.9% sensitivity and 93.75% specificity [11]. It has then been successfully used for identifying Rickettsiae in *Dermacentor marginalis* or *Rhipicephalus sanguineus* ticks [12]. For mosquito, it could distinguish *Plasmodium*-infected *Anopheles* mosquitoes from uninfected counterparts with high accuracy (98.75% in blind tests) [13], detect infection with filariae [14] or presence of *Wolbachia* in *Aedes* [15].

Identification of *Leishmania* species from cultures using MALDI-ToF is possible at least at complex level [16,17].

In a recent paper published de Andrade Silva et al. used MALDI-Tof to discriminate different sand flies’s peptide/protein profile according to sex, age and blood meal source [18]. This paper also showed 3 ions associated with *L. infantum* infection (m/Z 1984, 2010 and 2027). However, these 3 ions present a limited diagnostic performance with an AUC of 0.68, 95% CI [0.36, 0.99], in the model used for building datasets. Moreover, this paper lacks an external validation dataset, which is needed to assess the generalisation of the models [19].

Our objective is to identify MALDI-ToF protein signatures associated with the presence of *L. infantum* in *Lu. longipalpis* and *Ny. neivai* to propose a high-throughput infection detection method in these vectors.

## Material and methods

### Sand flies’ colonies

For the experimental assays, we used colonies of *Lu. longipalpis*, originally collected in Jacobina (Bahia, Brazil), and *Ny. neivai*, collected in Adrianópolis (Paraná, Brazil). Both colonies were maintained at the insectary of the Insect Vectors and Parasites Laboratory, Federal University of Paraná, following the rearing protocol described by Lawyer et al. (2017). Sand flies were reared in a climate-controlled chamber under standardized conditions (26 ± 1 °C; 80% relative humidity). Adult specimens were provided with cotton pads soaked in a 30% sugar solution as a food source.

### Experimental design

Promastigotes of *Leishmania infantum* (strain WHO/MHOM/BR/74/PP75) were cultured in RPMI 1640 medium supplemented with 10% heat-inactivated fetal bovine serum and 1% penicillin-streptomycin. Parasites in the logarithmic growth phase were harvested and mixed with heat-inactivated rabbit blood to a final concentration of 1 × 10⁷ parasites/mL. The infected blood was placed in glass feeders lined with chicken skin and offered to approximately 70 female *Lu. longipalpis* and *Ny. neivai* under controlled laboratory conditions. Fully engorged females were collected on days 2, 4, 5, 6, and 7 post-infection and immediately stored at –80 °C until analysis by MALDI-ToF mass spectrometry (Figure 1).

**Figure 1:**
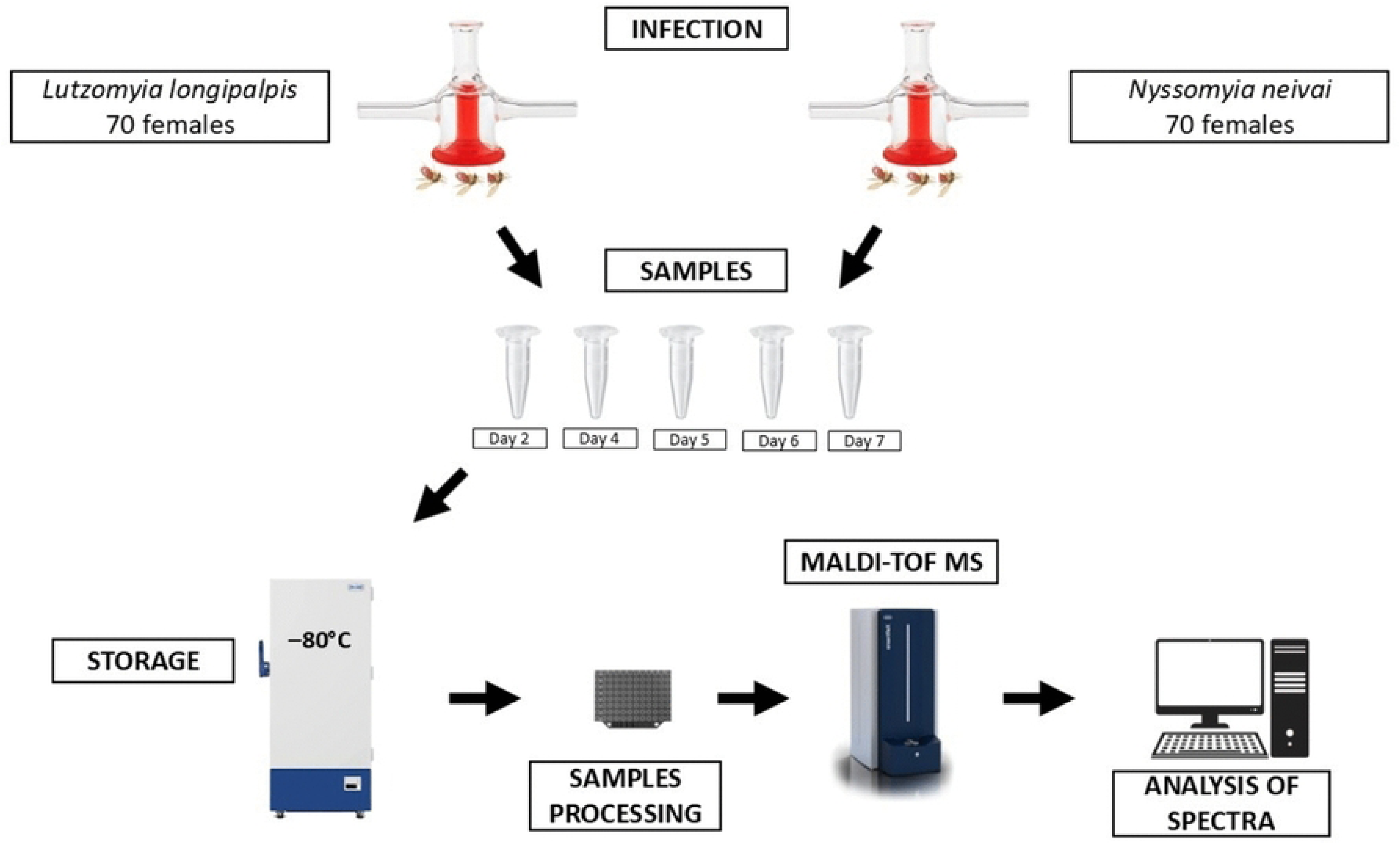
Schematic representation of the experimental design for the detection of *Leishmania infantum* in sand flies using MALDI-ToF MS. Female *Lutzomyia longipalpis* and *Nyssomyia neivai* (n = 70 per group) were subjected to an infective blood meal, while non-infected *Lu. longipalpis* females were used as a negative control. After blood feeding, sand flies were collected at different time points post-infection (D2, D4, D5, D6 and D7), where D followed by numbers indicates the days after infection. Samples were stored at -80 °C, processed, and analyzed by MALDI-ToF mass spectrometry, followed by spectral analysis.

### MALDI-ToF Spectrometry

We conducted MALDI-ToF MS as already described [20]. The thoraxes and legs of the specimens were placed in 10 µL of formic acid (Sigma-Aldrich, Lyon, France) and manually ground using a Teflon pestle [21]. Subsequently, 10 µL of acetonitrile (Sigma-Aldrich) was added. After a brief centrifugation at 10,000 rpm for 2 minutes, 1 µL of the supernatant was applied to a 96-well steel MALDI target plate (Bruker Daltonics, Champs-sur-Marne, France) in octuplets. Once dried, 1 µL of HCCA matrix solution (Bruker Daltonics) was added, and the plates were left to dry at room temperature. Spectra were acquired using a Bruker Sirius One MALDI-ToF spectrometer, with each well being measured at least eight times. Instrument calibration was performed using the Bacterial Test Standard from Bruker Daltonics.

### Data analysis and Machine Learning

Spectra were visually inspected with FlexAnalysis v3.4 and analyzed using Biotyper Compass Explorer v4.1.100. The “MALDI Biotyper Preprocessing Standard Method” from Bruker was employed to create main spectra profiles (MSP). Hierarchical cluster analysis MSP dendrogram (HCA) was performed on MSP using MALDI-Biotyper Compass Explorer v4.1 software, with the correlation method for calculating distances and Ward algorithm for clustering.

Bruker FID spectra files were imported in R v4.5.1 [22] with the MALDIQuantForeign package [23]. High-quality spectra were preprocessed and selected using the screenSpectra function from the MALDIrppa package [24] using a 1.5 A-score threshold. Spectra were trimmed for the 3000-15000 region, smoothed using Savitzky-Golay algorithm. Baseline was removed using SNIP methods and intensities were calibrated based on total ion current. Peaks were detected using detectPeaks MALDIQuant function with a signal to noise ratio of 3 and a half window size of 20 and a tolerance of 0.005 for peaks binning. Peaks were aligned using alignPeaks from the MALDIrppa package. Spectral data were converted in an intensity peaks matrix from which datasets were built.

We performed a Principal Component Analysis (PCA) on the complete dataset and on subsetted datasets organised by the number of days after infection, using the R packages FactoMineR and factoextra [25,26].

Custom R scripts utilising the caret package were used for machine learning discrimination between infected and non-infected. [27]. The dataset used to build and validate the models was divided into training/validation (in an 80/20 proportion). An external “test dataset”, constituted of spectra from specimens not included in the training and validation datasets, was used to evaluate the performance of the models in a less biased situation. Predictors with unique variance were eliminated with the nearZeroVar function. To deal with class imbalance in the training/validation datasets, we used the down-sampling function of the caret package, subsetting the majority class to obtain frequency matching the least prevalent class. Five widely used machine-learning algorithms were used within the caret package: Support Vector Machine (SVM) with linear kernel, k-nearest neighbor (KNN), partial least squares discriminant analysis (PLS), Shrinkage discriminant analysis (SDA) and random forest (RF). Models were trained with repeated cross validation (ten folds and ten repeats). Performances during model training were evaluated based on ROC metrics with the twoClassSummary function. Relevant parameters tuning was performed using a tuning grid. Final models’ performances were evaluated on the validation dataset and the external “test dataset” based on accuracy, sensitivity and specificity. Variable importance was evaluated for each model using the varImp function of caret package. Venn diagrams were plotted using the ggVennDiagram package [28].

## Results

### Acquisition of spectra

Female adult specimens of *Lu. longipalpis* were processed individually (n=50), 26 specimens were not infected and 24 infected. After elimination of flat or bad quality spectra, 3038 spectra remained. We obtained 1588 spectra from non infected females and 1450 from infected females. The number of collected spectra according to condition is described in Supplementary Table S1. Representative spectra of infected and not infected specimens at D2, D4 and D6 are presented in Figure 2 (panel A and B). Despite variability across age and infection status, spectra showed several constant peaks complexes (around m/z 4675, 5286, 6257, 8865, 9350).

**Figure 2:**
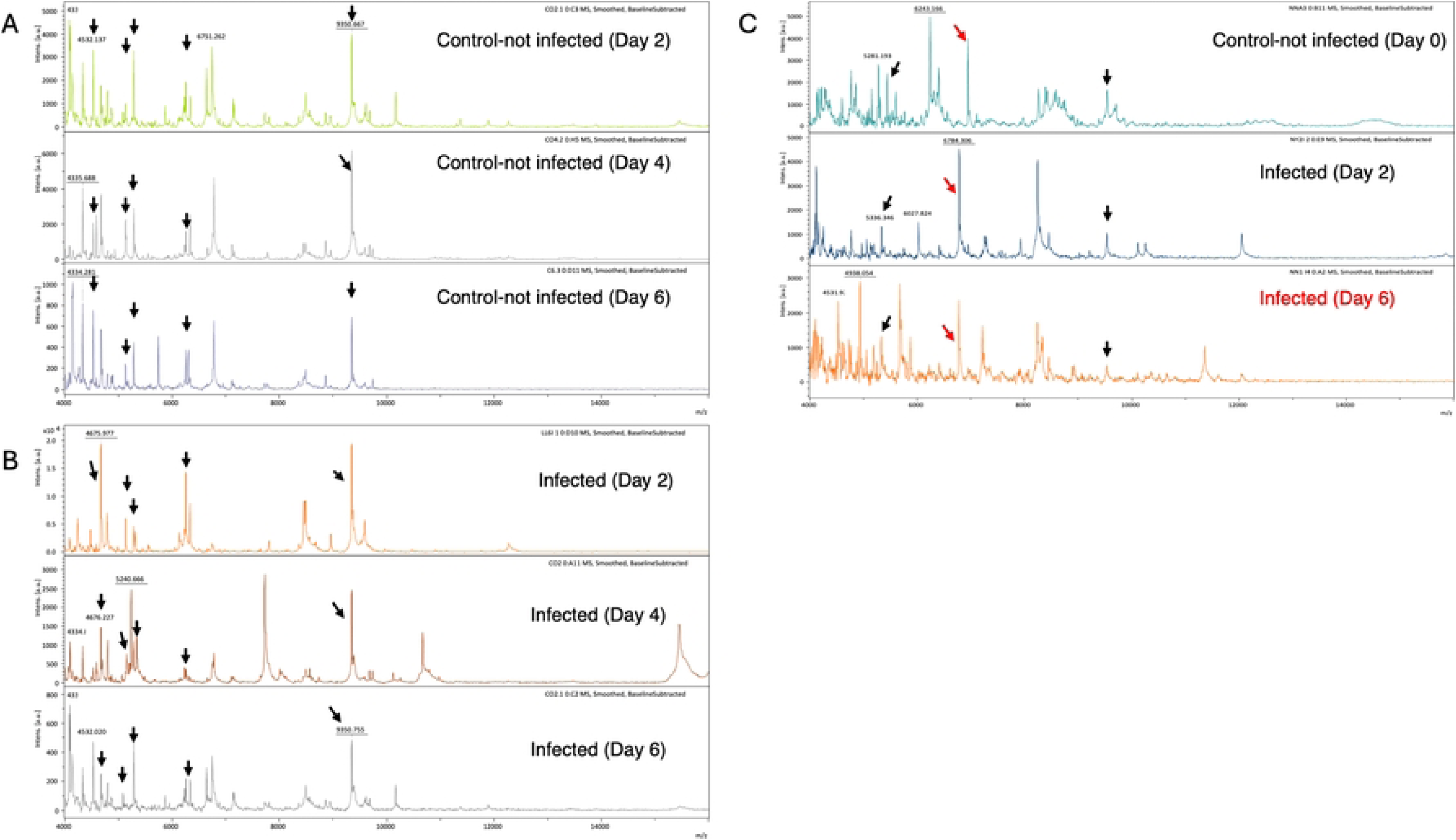
Representative spectra of A*) Lutzomyia longipalpis* not infected at D2, D4, D6, B) *Lutzomyia longipalpis* infected at D2, D4, D6. the most visible constant peak complexes are indicated by black arrows. C) *Nyssomyia neivai* not infected at D0 and infected at D2 and D6, most visible constant peak complexes are indicated by black arrows and peak shift around 6784 is indicated by red arrow.

Female adult specimens of *Ny. neivai* were processed individually (n=22), 6 specimens were not infected and 16 infected. After elimination of flat or bad quality spectra, 1277 spectra remained. We obtained 355 spectra from non infected females and 922 from infected females. Due to the difficulty associated with breeding of *Ny. neivai*, all spectra from non-infected samples were from specimens collected at D0 (6 specimens, 355 spectra). The number of collected spectra according to condition is described in Supplementary Table S1. Representative spectra of non-infected (D0) and infected specimens at D2 and D6 are presented in Figure 2 (panel C). Visual inspection of spectra showed several constant peaks complexes (for example around m/z 5336 and 9540) and some peak-shifts (for example from 6949 in non-infected to 6781-6784 in infected).

### Unsupervised classification of infected and non-infected specimens of Lu. longipalpis and Ny. neivai

Hierarchical classification analysis of the MSPs from *Lu. longipalpis* spectra showed a grouping into two clusters (Figure 3 A). Cluster A included 12 MSPs from infected specimens (from D2 to D7) and one MSP from a non-infected specimen on D2. Cluster B included spectra from 14 infected specimens (from D2 to D7) and ten MSPs from non-infected specimens (from D4 to D6). The distance between MSPs from infected specimens and MSPs from uninfected specimens in the different sub-clusters of the tree was small (> 200). Principal component analysis of the complete datasets did not reveal discriminant components able to distinguish infected from non-infected samples (Figure 4 panel A). However, when performed in datasets subsetted by day from experimental infection, the first two components (ie the two components explaining the largest amount of variance) permitted clustering of individuals according to their infected or not infected status (Figure 4 panel B-F).

**Figure 3.**
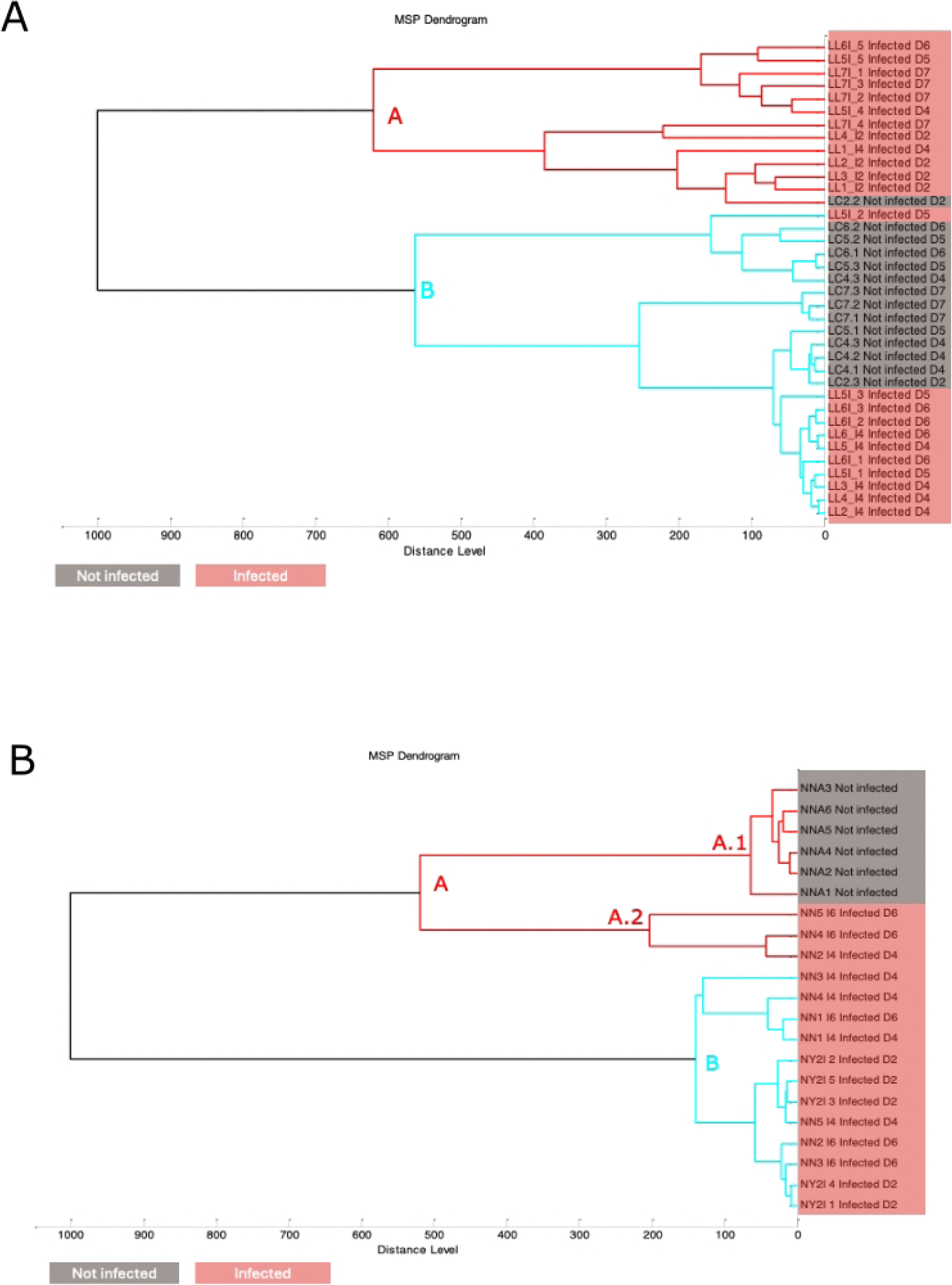
MSP dendrogram of Infected and not infected spectra at various days post infection A) *Lutzomyia longipalpis*, B) *Nyssomyia neivai*

**Figure 4:**
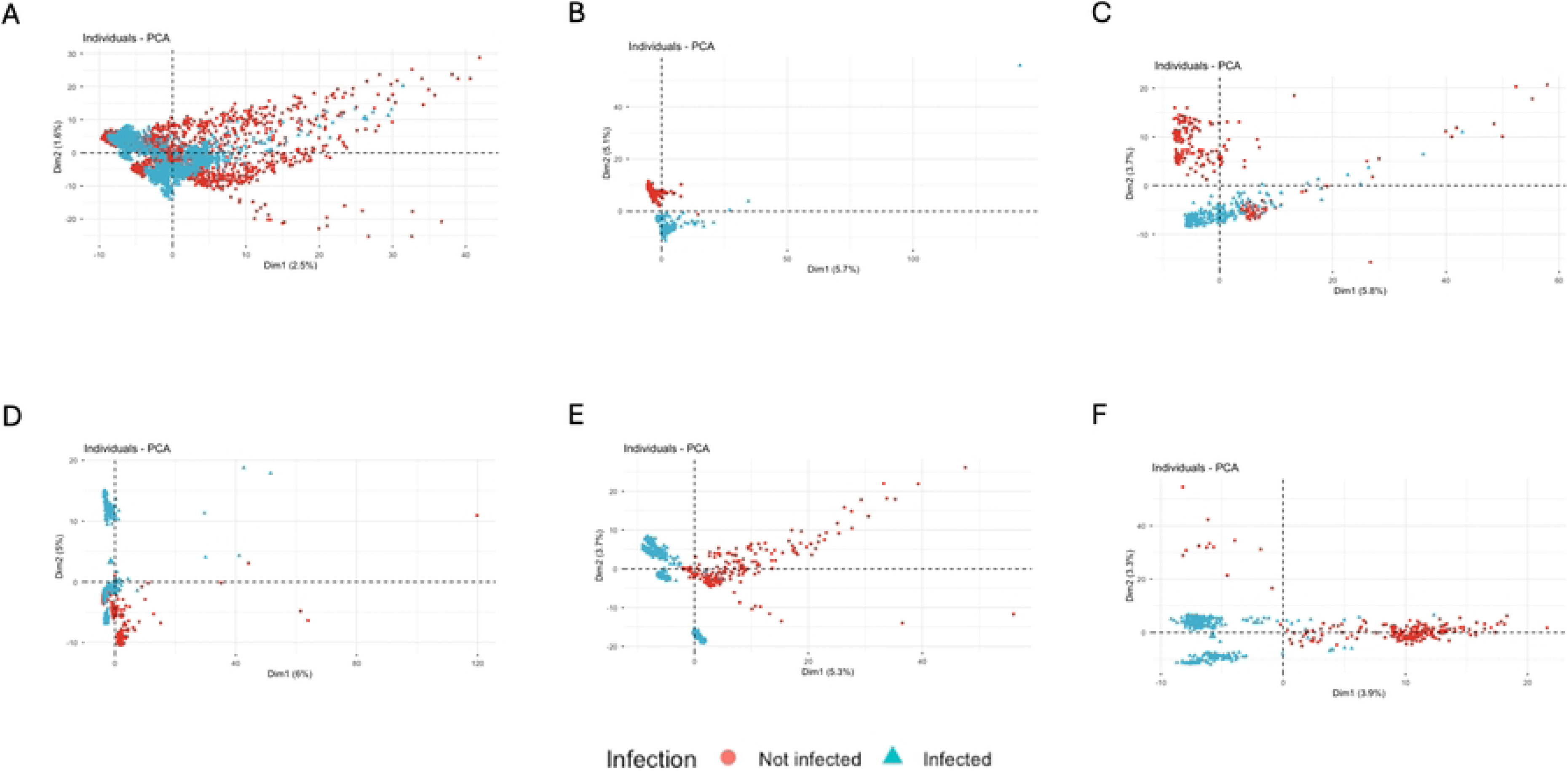
Principal component analysis of *Lutzomyia longipalpis* according to infection status. Biplot of the two first component (respectively Dim1 and Dim2) A) whole dataset, B) Day 2 C) Day 4 D) Day 5 E) Day 6 and F) Day 7

Concerning spectra acquired from *Ny. neivai* specimens, hierarchical classification analysis of the MSPs showed a grouping into two clusters (Figure 3B). Cluster A included 7 MSP from non-infected specimens (subcluster A.1) and 3 MSPs from infected specimens (from D4 and D6, subcluster A.2). Cluster B included spectra from 12 infected specimens (from D2 to D6). The distance between sub-clusters from infected specimens and sub-clusters from uninfected specimens was important (> 500 in cluster A). Principal component analysis of the complete datasets revealed good discrimination of infected from non-infected samples with components 1 and 2 (Figure 5 panel A). Analysis by day from experimental infection on the first two components showed the consistent distinct clustering of specimens at D0 (not infected) and specimens at D7. The other specimens were more dispersed. The uninfected and D7 clusters were closest according to dimension 1 (Figure 5 panel B).

**Figure 5:**
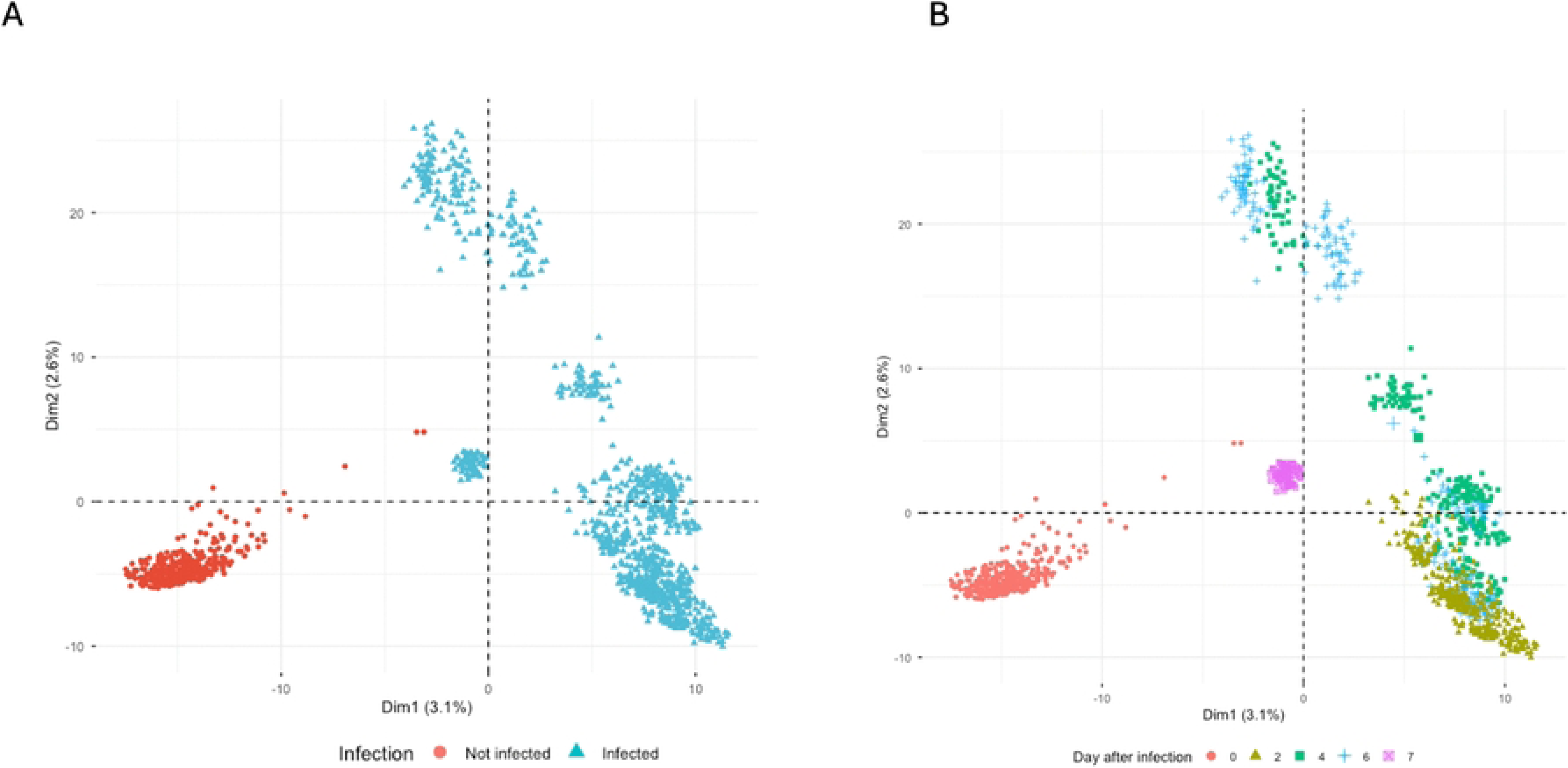
Principal component analysis of *Nyssomyia neivai* according to infection status. Biplot of the two first component (respectively Dim1 and Dim2) A) whole dataset, infected specimen’s vs not-infected B) the whole dataset, specimens according to sampling day after infection, specimens not infected are labelled as Day after infection 0.

### Development of machine learning models for classification of infected and non-infected specimens of Lu. longipalpis and Ny. neivai

For development of machine-learning models for detection of *Lu. longipalpis* infection by *L. infantum*, the spectra from two infected (LAF4 at D0, and LC5.2 at D5) and two non infected specimens (LL1_I2 at D2 and LL5I_4 at D5) were assigned to the testing dataset. These spectra (119 non infected and 128 infected) were not used for building or validations of predictive models but only for validations of models. All others were used for training (n=977 for each class) and validation (n=244 for each class) of the different models. The complete dataset contained the intensity of 2005 peaks. No predictors had only one distinct value (unique variance). Results of the different machine learning algorithms on the test and validation dataset are given in Table 1, results are given in numbers of spectra. Except for SVM and SDA (1 misclassified spectra), all algorithms achieve perfect classification on the validation dataset. Excellent classification (>95% accuracy) was obtained on the external test for PLS, SDA, KNN and RF. Only one spectrum was misclassified on the test dataset with RF (Accuracy 99.6%, sensitivity 100%, specificity 99.16%). Using SVM, most spectra from the test dataset were classified as not-infected (Accuracy 44.5%, Sensitivity 46.9% and specificity 87.4%).

**Table 1:** performance of the machine learning algorithm in validation and test datasets, both for Lu. longipalpis and Ny. neivai. TP: True positive, FP: False positive, FN: False negative, TN: True negative.

The average spectra of non-infected samples (Figure 6, panel A) from the training/validation dataset revealed a characteristic proteomic baseline across the m/z range of 2,000–20,000. In contrast, spectra from infected specimens (Figure 6, panel B) exhibited marked intensity elevations or decreases at multiple m/z regions, reflecting infection-associated proteomic remodelling. The variable importance dot plot of the three top-performing models (PLS-DA, Random Forest and SDA) highlighted regions of interest for infection biomarkers (Figure 6, panel C). Interestingly, the important variables differed according to the model. To determine the important variables shared or not by the models, we used Venn diagram analysis to compare the m/z features that exceeded the 1% importance threshold in PLS, RF and SDA (Figure 6, panel D). This analysis revealed that only 219 peaks (45% of the total) were shared by all three models, with an additional 5 (1%) PLS-specific, 175 (36%) SDA-unique and 14 (3%) RF-distinctive markers. Concerning the two others models (Figure 6, panel E), SVM and KNN shared 238 peaks (50%) with PLS and 220 (46%) betwen them. A comparison of the top three most important variable across the 3 models (m/Z 4239, 4331, 5245, 6783, 7736, 8489, 9562 and 12279) demonstrate clear separation by infection status, with infected specimens showing significantly decreased (m/Z 4331,5245,6783,7736) or increased (m/Z 8489,9562,12279) intensities versus non-infected controls (p-adjusted < 0.05 using Bonferroni correction, t-test) (Figure 6, panel F). Comparing the intensity of these six peaks across different days post-infection showed variability over time. (Supplementary Figure 1).

**Figure 6:**
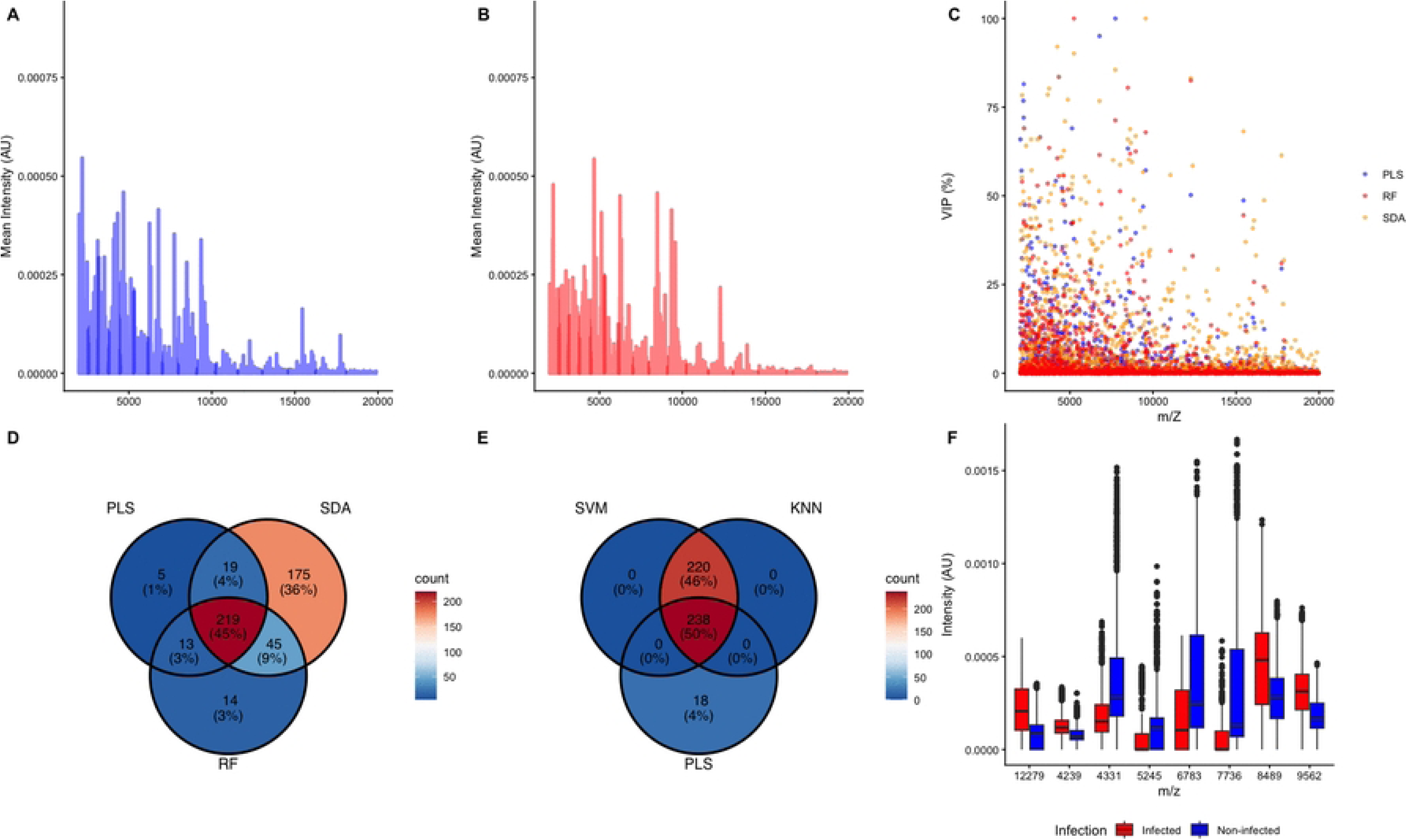
Variable used for model prediction of infection in *Lu. longipalpis*. A) Average spectra of non-infected specimens B) Average spectra of infected specimens C) Dot-plot of variable importance for each m/Z value for the three best models (PLS, RF, SDA) D) Venn Diagram of the variable with importance >1% used by the 3 models (PLS, RF, SDA) E) Venn Diagram of the variable with importance >1% used by the 3 models (PLS, SVM, KNN) F)Boxplot of the top 3 variable in the three best model, according to infection status.

We also developed machine-learning models for detecting *L. infantum* infection in *Ny. neivai*. Spectra from two non-infected specimens (NNA4 and NNA5) and two infected (NY2I at D2 and NN1_I6 at D6) were assigned to the testing dataset. These spectra (121 non infected and 116 infected) were not used for building or validations of predictive models. Remaining specimens were used for training (n=192 for each class) and validation (n=48 for each class) of the different models.

The eight algorithms used showed high performance to distinguish infected from non-infected females of *Ny. neivai* with accuracy, sensitivity and specificity superior to 99% [Table 1]. Only one false-positive was observed with SVM, KNN, PLS and RF in the external testing dataset. The average spectra of non-infected samples (Figure 7, panel A) from the training/validation dataset revealed a characteristic proteomic baseline across the m/z range of 2,000–20,000. In contrast, spectra from infected specimens (Figure 7, panel B) exhibited marked intensity elevations or decreases at multiple m/z regions, reflecting infection-associated proteomic remodelling. The variable importance dot plot of three models (PLS-DA, Random Forest and SDA) highlighted regions of interest for infection biomarkers (Figure 7, panel C and). Interestingly 217 important variables (42%) for classification were common between SDA, PLS and RF. More than half of the variables of interest (283 representing 56%) were common between PLS and SDA. A comparison of the top three most important variable of RF, PLS and SDA models (m/Z 2984, 4090, 5190, 6246, 6315, 6486, 6783 and 8549) demonstrate variations by infection status, with infected specimens showing significantly increased (m/Z 2984, 4090, 5190, 6783) or decreased (m/Z 6246, 6315, 6486, 8549) intensities versus non-infected controls (p-adjusted < 0.05 using Bonferroni correction, t-test) across all top biomarkers (Figure 7, panel F). Comparing the intensity of these six peaks across different days post-infection revealed variability over time in *Ny. neivai* (Supplementary Figure 2), similar to that observed with *Lu. Longipalpis* (Supplementary Figure 1).

**Figure 7:**
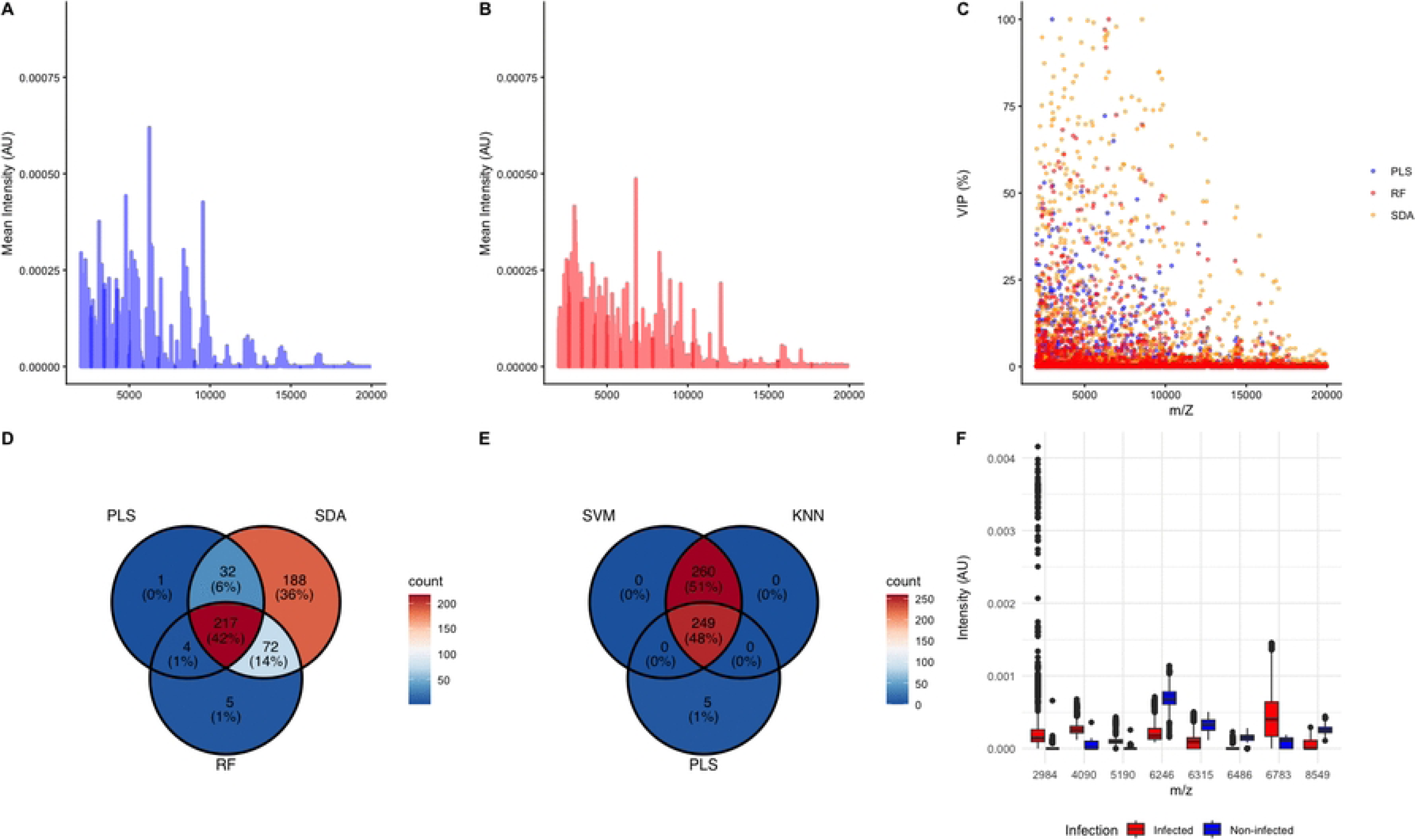
Variable used for model prediction of infection in *Ny. neivai*. A) Average spectra of non-infected specimens B) Average spectra of infected specimens C) Dot-plot of variable importance for each m/Z value for the three best models (PLS, RF, SDA) D) Venn Diagram of the variable with importance >1% used by the 3 models (PLS, RF, SDA) E) Venn Diagram of the variable with importance >1% used by the 3 models (PLS, SVM, KNN) F)Boxplot of the top 3 variable in the three best model, according to infection status.

Across the three best classification models (PLS, SDA and RF), the MALDI-ToF peak sets used to discriminate *Leishmania*-infected samples in *Lu. longipalpis* and *Ny. neivai* showed moderate overlap (using an importance threshold of 20%). In the PLS model, most peaks were vector-specific, with only 6 (5%) shared peaks between the two vectors (Figure 8, panel A), whereas SDA and KNN retained a larger common subset, with 59 (12%) shared peaks (Figure 8, panel C and D). The RF model also present a relatively limited overlap, with 41 (10%) peaks shared between the two species (Figure 8, panel B). Overall, low correlation and no linear relation was observed between the peak’s importance of the two species (Pearson correlation coefficient of 0.1724 and R2 0.0294 for the PLS model, Supplementary Figure 3).

**Figure 8:**
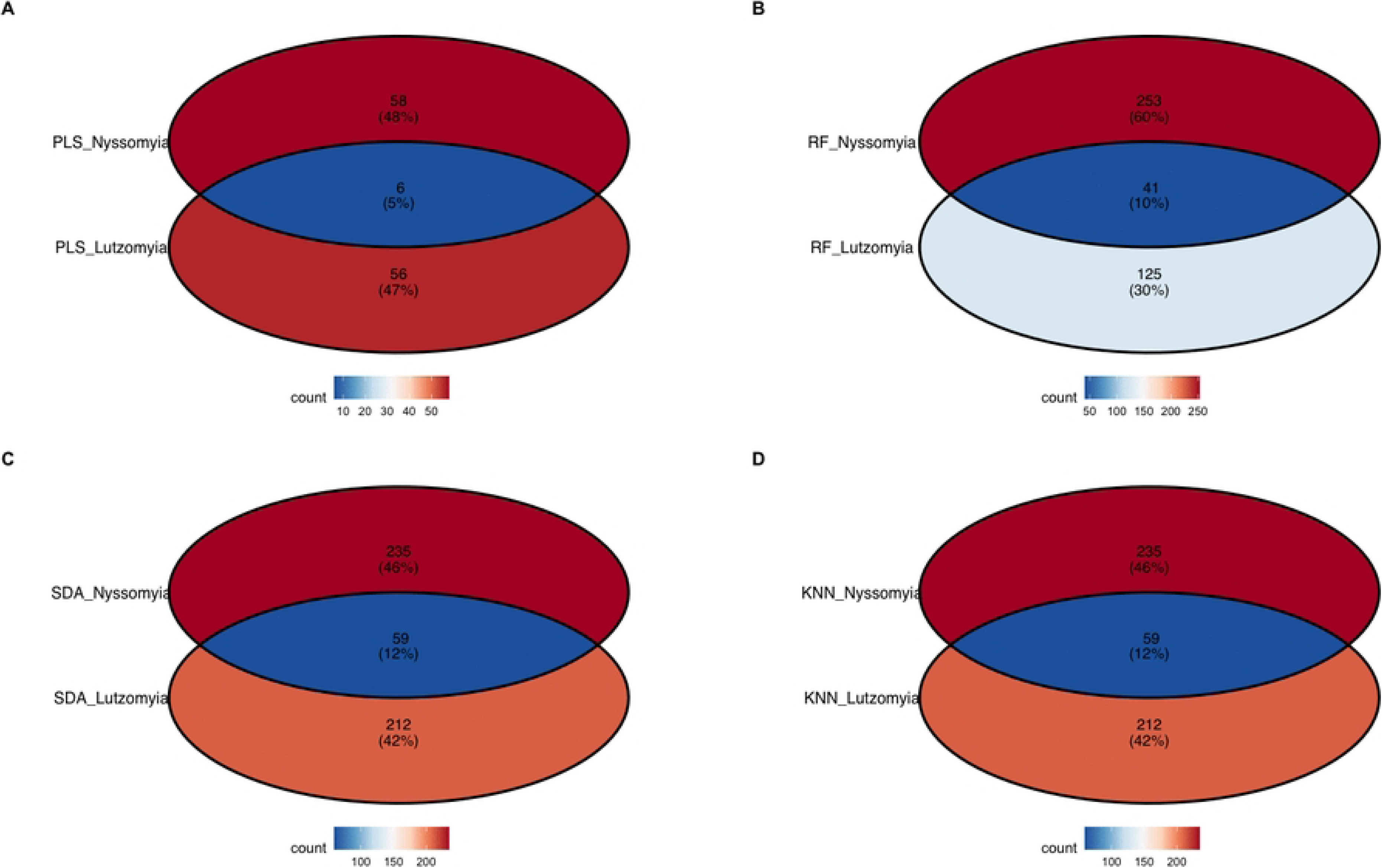
Variable used for model prediction with importance > 20. Venn Diagram of the intersection peaks between *Lu. longipalpis* and *Ny. neivai* with four classification models. A) PLS B) RF C) SDA D) KNN Numbers indicate the count of selected peaks in each vector-specific region and in the overlap. Percentages show the proportion of peaks in each subset relative to the total selected within each model.

## Discussion

### Main findings

This pilot study shows that *L. infantum* infection is associated with reproducible changes in the MALDI-TOF MS protein profiles of *Lu. longipalpis* and *Ny. neivai* females. Unsupervised analysis (MSP clustering and PCA on complete datasets) provided limited or inconsistent discrimination between infected and uninfected specimens, in the case of *Lu. longipalpis*. Indeed, low distance in HCA between infected and non-infected specimens indicate a limited ability of distance-based techniques to discriminate between spectra for *Lu. longipalpis*. For *Ny. neivai*, the high distance between infected and non-infected MSP from infected and not infected specimens indicate a potential ability of distance-based techniques to discriminate between spectra from infected specimens and spectra from uninfected specimens. This high distance could also possibly be a bias explained by the absence of not infected samples after day 0.

Supervised machine-learning models trained on peak-intensity matrices have been shown to achieve excellent classification performance in both species. To limit the possibility of bias due to overfitting, we confirmed these results using an external test set composed of spectra from individuals who were not used for model training or internal validation.

For *Lu. longipalpis*, several algorithms (PLS-DA, SDA, RF, KNN) correctly classified almost all spectra, in the test external dataset, with only linear SVM showing residual misclassification. For *N. neivai*, all algorithms reached high accuracy, sensitivity and specificity >99%, with only a single false-positive prediction using SDA.

Variable-importance analyses revealed sets of discriminant m/z features, many of which decreased in intensity in infected sand flies, while a few increased, suggesting infection-associated proteomic remodelling rather than simply the appearance of parasite-specific peaks. The list of candidate biomarkers differs partly between models and between species, but some peaks were consistently highlighted across algorithms, especially for *Ny. neivai*. Collectively, these results indicate that MALDI-ToF MS, combined with supervised learning, can detect subtle infection-related changes in sand-fly protein fingerprints, even when unsupervised methods fail to separate groups.

### Comparison with previous work

Our results extend previous applications of MALDI-ToF MS to sand flies and *Leishmania*. Earlier studies focused mainly on species identification using legs or whole flies and demonstrated high accuracy for Neotropical *Lutzomyia* and *Nyssomyia* species [10]. More recently, de Andrade Silva et al. showed that peptide/protein profiles of *Lu. longipalpis* vary according to sex, age, blood meal source and *L. infantum* infection, identifying three ions associated with infection (m/z 1984, 2010 and 2027), but with modest diagnostic performance (AUC 0.68, wide confidence interval) and without external validation [18].

The present study confirms that infection alters sand-fly MALDI-TOF profiles but differs in three important aspects. First, we analysed two vector species, *Lu. longipalpis* and *Ny. neivai*, which enabled us to show that infection-related signatures are at least partly species-specific and that *Ny. neivai* exhibits particularly clear separation between infected and uninfected profiles. Second, we used a dedicated external test dataset composed of spectra from individuals not used in training, providing a more robust assessment of model generalisability than internal cross-validation alone. Third, instead of focusing only on the most intense peaks, we used full peak-intensity matrices and compared different algorithms, which revealed that informative biomarkers are not restricted to the highest-intensity ions and that different models rely on overlapping but non-identical sets of peaks.

These observations are consistent with prior work on MALDI-TOF detection of pathogens in other vectors, such as *Rickettsia*-infected ticks or *Plasmodium*-infected mosquitoes [12], where supervised models can accurately classify infection status even when infection-related variance is not captured by the first principal components.

### Biological interpretation and alternative explanations

The candidate biomarkers identified here likely reflect a complex mixture of parasite-derived proteins, vector proteins modulated by infection, and possibly host-derived components (e.g. residual blood or immune factors) that persist in the thorax and legs. Therefore, it is not yet possible to attribute individual peaks to parasite versus vector origin. The spectra probably integrate a lot of proteins from vector muscle, cuticle, salivary glands, haemolymph and residual midgut content. Differences between infected and uninfected profiles could therefore reflect systemic immune responses, altered feeding status or tissue damage, not just the presence of parasite proteins. The predominance of peaks with decreased intensity in infected specimens suggests that infection may down-regulate or degrade specific sand-fly proteins, or alter their ionisation efficiency, rather than simply adding peaks from parasite proteins or from the vector immune response. This is compatible with previous studies reporting remodelling of sand-fly salivary and midgut proteins during *Leishmania* infection, even if this response seem limited [29]. Interestingly the different stages of *Leishmania* development was associated to specific transcriptional variation, which could explain the changes in protein profiles observed by MALDI-ToF over time [30]. Vector immune responses and tissue damage associated with infection can also substantially change protein expression patterns [31,32]. These findings could also show that future works combining MALDI-ToF with compartment-specific dissections (e.g. isolated salivary glands or midguts) and targeted proteomics will be needed to clarify the biological origin of these discriminant peaks. An alternative approach would be to use other proteomic tools, such as LC-MS/MS, to identify the proteins corresponding to the different biomarkers. These could then be compared to predicted or identified proteins from vectors and *Leishmania* genomes.

### Methodological limitations

A major limitation of this study is the absence of direct confirmation of infection status in each individual sandfly by dissection or PCR. Our classification of specimens as “infected” or “non-infected” relied on experimental infection design and colony controls, but infection could not be verified in the exact individuals used for MALDI-ToF. This choice was deliberate: as a proof-of-concept, we prioritised preserving all potentially infected specimens for spectral analysis and avoided dissection that would have destroyed material. The same limitation affects the study by de Andrade Silva et al., indicating that this is a broader challenge in MALDI-ToF–based infection studies. Nevertheless, previous experimental infections of *Lu. longipalpis* with *L. infantum* have reported relatively high infection rates under laboratory conditions, often ranging from approximately 40% to 80% depending on parasite dose, feeding conditions and parasite strain (Freitas et al., 2012; Sant’Anna et al., 2014; Alexandre et al., 2020). These studies indicate that a substantial proportion of females typically become infected following artificial blood feeding, supporting the assumption that a significant fraction of the experimentally exposed sand flies used here were indeed infected. Our classifiers also did not estimate infection intensity nor discriminate parasite-derived from vector-derived peaks.

In future experiments, we plan to combine MALDI-ToF and microscopic/molecular diagnostics on the same individual. This will allow direct quantification of sensitivity and specificity at the specimen level, facilitate the calibration of machine-learning models against true infection status, and help disentangle effects of infection intensity or stage on the spectra.

Finally, the sample size, particularly for *Ny. neivai*, remains modest, and the experimental infection setting may not fully reproduce the heterogeneity of field populations in terms of age structure, microbiota, and co-infections. External validation on independent colonies and, ultimately, on field-caught sand flies from endemic areas will be essential before MALDI-ToF-based infection classification can be proposed for operational surveillance.

## Conclusion

Despite these limitations, this study demonstrates that MALDI-ToF MS, combined with appropriate data processing and supervised learning, can detect *L. infantum* infection in *Lu. longipalpis* and *Ny. neivai* with high accuracy under the experimental conditions. In the longer term, such an approach could be integrated into entomological surveillance as a high-throughput tool to screen large numbers of vectors for both species identification and infection status, for a very low cost. This would be particularly attractive in settings where MALDI-ToF is already deployed for clinical microbiology.

For field application, several steps are needed: optimisation of specimen preservation and transport compatible with MALDI-ToF analysis, validation of models on wild populations with natural infection loads, and integration of MALDI-ToF data with PCR-based detection of *Leishmania* and host-blood origin in the same individuals. If these challenges can be addressed, MALDI-ToF-based detection of infected sand flies could complement existing surveillance tools and provide a cost-effective means to monitor transmission risk in visceral leishmaniasis foci where *Lu. longipalpis* and *Ny. neivai* coexist or where *Ny. neivai* may act as a secondary vector.

## Acknowledgements

We would like to thank all the staff at UR ESCAPE (University of Reims Champagne-Ardenne) and the Insects, Vectors, and Parasites Laboratory at the Federal University of Paraná who were involved in maintaining the sandfly colonies and collecting data.

## CRediT authorship contribution statement

Conceptualization: LAS, JD, MCC, AH; Data curation: LAS, EK, AH; Formal Analysis: LAS, AH; Funding acquisition: JD, MCC; Investigation: LAS, EK, JP; Methodology: LAS, EK, JP, AH; Project administration: LAS, JP, JD, MCC, AH; Resources: JD, MCC; Software: AH; Supervision: JD, MCC, AH; Validation: LAS, EK, JP, JD, MCC, AH; Visualization: LAS, EK, AH; Writing – original draft: LAS, JD, AH; Writing – review & editing: LAS, EK, JP, JD, MCC, AH

## Data availability

For reproducibility purposes under Findability, Accessibility, Interoperability, and Reusability (FAIR) principles, raw spectra and R scripts used for the analyses developed in this study are provided. Raw spectra were deposited in Zenodo under doi: https://doi.org/10.5281/zenodo.20699798. R scripts are available from Github: https://github.com/Ornithodoros/sandfly_leishmania_MALDI_detection

---

Supplementary figure 1: Variation of the top 3 variable in the three best model (PLS, SDA, RF), according to the infection status and the number of day post-infection. in *Lu. longipalpis* A) Not infected specimens B) Infected specimens

Supplementary figure 2: Variation of the top 3 variable in the three best model (PLS, SDA, RF), according to the infection status and the number of day post-infection. in Ny. neivai A) Not infected specimens B) Infected specimens

Supplementary figure 3: Heatmap and dot-plot comparing importance scores of variables used by PLS model, both in *Lu. longipalpis* and *Ny. neivai*.

**Table S1:** Number of specimens and spectra according to conditions.

| infection status | Age in day after infection | number of spectra having pass QC | number of specimens |
| --- | --- | --- | --- |
| *Lutzomyia longipalpis* |  |  |  |
| Not infected | 0 | 431 | 8 |
| Not infected | 2 | 186 | 3 |
| Not infected | 3 | 191 | 3 |
| Not infected | 4 | 225 | 3 |
| Not infected | 5 | 193 | 3 |
| Not infected | 6 | 183 | 3 |
| Not infected | 7 | 179 | 3 |
| Infected | 2 | 298 | 5 |
| Infected | 4 | 282 | 5 |
| Infected | 5 | 318 | 5 |
| Infected | 6 | 310 | 5 |
| Infected | 7 | 242 | 4 |
| *Nyssomyia neivai* |  |  |  |
| Not infected | 0 | 355 | 6 |
| Infected | 2 | 303 | 5 |
| Infected | 4 | 278 | 5 |
| Infected | 6 | 277 | 5 |
| Infected | 7 | 64 | 1 |

